# Inflammation and neuronal gene expression changes differ in early vs late chronic traumatic encephalopathy brain

**DOI:** 10.1101/2022.05.31.494179

**Authors:** Adam Labadorf, Filisia Agus, Nurgul Aytan, Jonathan Cherry, Jesse Mez, Ann McKee, Thor D. Stein

## Abstract

Our understanding of the molecular underpinnings of chronic traumatic encephalopathy (CTE) and its associated pathology in post-mortem brain is incomplete. Factors including years of play and genetic risk variants influence the extent of tau pathology associated with disease expression, but how these factors affect gene expression, and whether those effects are consistent across the development of disease, is unknown. To address these questions, we conducted an analysis of the largest mRNASeq whole-transcriptome dataset available to date. We examined the genes and biological processes associated with disease by comparing individuals with CTE with control individuals with a history of repetitive head impacts that lack CTE pathology. We then identified genes and biological processes associated with total years of play as a measure of exposure, amount of tau pathology present at time of death, and the presence of *APOE* and *TMEM106B* risk variants. Samples were stratified into low and high pathology groups based on extent of tau pathology and years of play to model early vs late changes in response to exposure, and the relative effects associated with these factors were compared between these groups. Substantial gene expression changes were associated with severe disease for most of these factors, primarily implicating diverse, highly increased neuroinflammatory and neuroimmune processes. In contrast, low exposure groups had many fewer genes and processes implicated and show striking differences for some factors when compared with severe disease. Specifically, gene expression associated with amount of tau pathology showed a nearly perfect inverse relationship when compared between these two groups. Together, these results suggest the early disease process may differ substantially from that observed in late stages, that total years of play and tau pathology influence disease expression differently, and that related pathology-modifying risk variants may do so via distinct biological pathways.

## 1 Introduction

Chronic traumatic encephalopathy (CTE) is a progressive neurodegenerative disease associated with prolonged exposure to traumatic head impacts most commonly observed in the brains of athletes of contact sports and combat veterans. CTE pathology is strongly associated with other neurological disorders, including depression, dementia, Alzheimer’s and Parkinson’s disease, and clinical symptoms include behavioral, personality and mood changes, and memory and cognitive function deficits. Upon autopsy, CTE is diagnosed by the presence of neurofibrillary tangles found around blood vessels in the depths of the cortical sulcus.

The primary risk factor for developing CTE is the duration of exposure to repetitive head impacts, which is measured in total years of play for athletes^1^. The neurological symptoms of CTE often only manifest decades after players have retired, suggesting the effects of repetitive head impacts set in motion a progression of pathological events ultimately leading to disease. However, not all individuals with a similar degree of exposure will go on to develop CTE, and while some genetic evidence suggests specific genes are associated with increased risk of disease, the mechanisms underlying the disease process are currently unknown. It is also currently unknown whether early disease states differ from those that are coincident with severe pathology.

Our previous work showed that two genetic risk variants in the Apolipoprotein E (APOE) and Transmembrane Protein 106B (TMEM106B) genes also influence the frequency and severity of CTE. The association of the *APOE e4* allele with increased Alzheimer’s Disease (AD) risk is well documented^2^ and *APOE e4* has also been shown to be associated with increased severity of CTE pathology (internal discussions with BU CTE Center investigators, findings in submission at time of writing). Variants in *TMEM106B* are associated with increased neuroinflammation in aging^3^, frontotemporal lobar degeneration (FTLD)-TDP^4^, and with AD^5,6^, and our prior study suggested a protective role of the TMEM106B variant rs3173615 in CTE^7^. While prior research has provided some information about the functional roles of APOE^8^ and to a lesser extent TMEM106B^9–12^ in both normal and disease contexts, the role they play in the development of CTE is poorly understood.

To address these knowledge gaps, we profiled whole transcriptome gene expression of prefrontal cortex (Brodmann Area 9) by mRNA-Seq in a cohort of 66 CTE and 10 repetitive head impacts (RHI) controls donated to the VA-BU-CLF Brain Bank. The CTE samples were subdivided into 13 low (CTE-L, Stages I & II) and 53 high (CTE-H, Stages III & IV) pathology groups using the McKee staging criteria^13,14^, with the goal of identifying molecular changes associated with early versus late disease. The RHI controls are individuals who experienced a similar level of repetitive head impacts exposure as the disease groups but displayed no CTE pathology upon autopsy. In this study, we sought to identify genes and biological processes associated with differences between CTE and RHI controls, repetitive head impacts exposure as measured by total years of play, amount of tau pathology using immunohistological quantification of Phospho-Tau (AT8), and between APOE and TMEM106B risk variant carrier groups.

## 2 Results

Distributions of these key sample characteristics are depicted in Figure 1.

**Figure 1.**
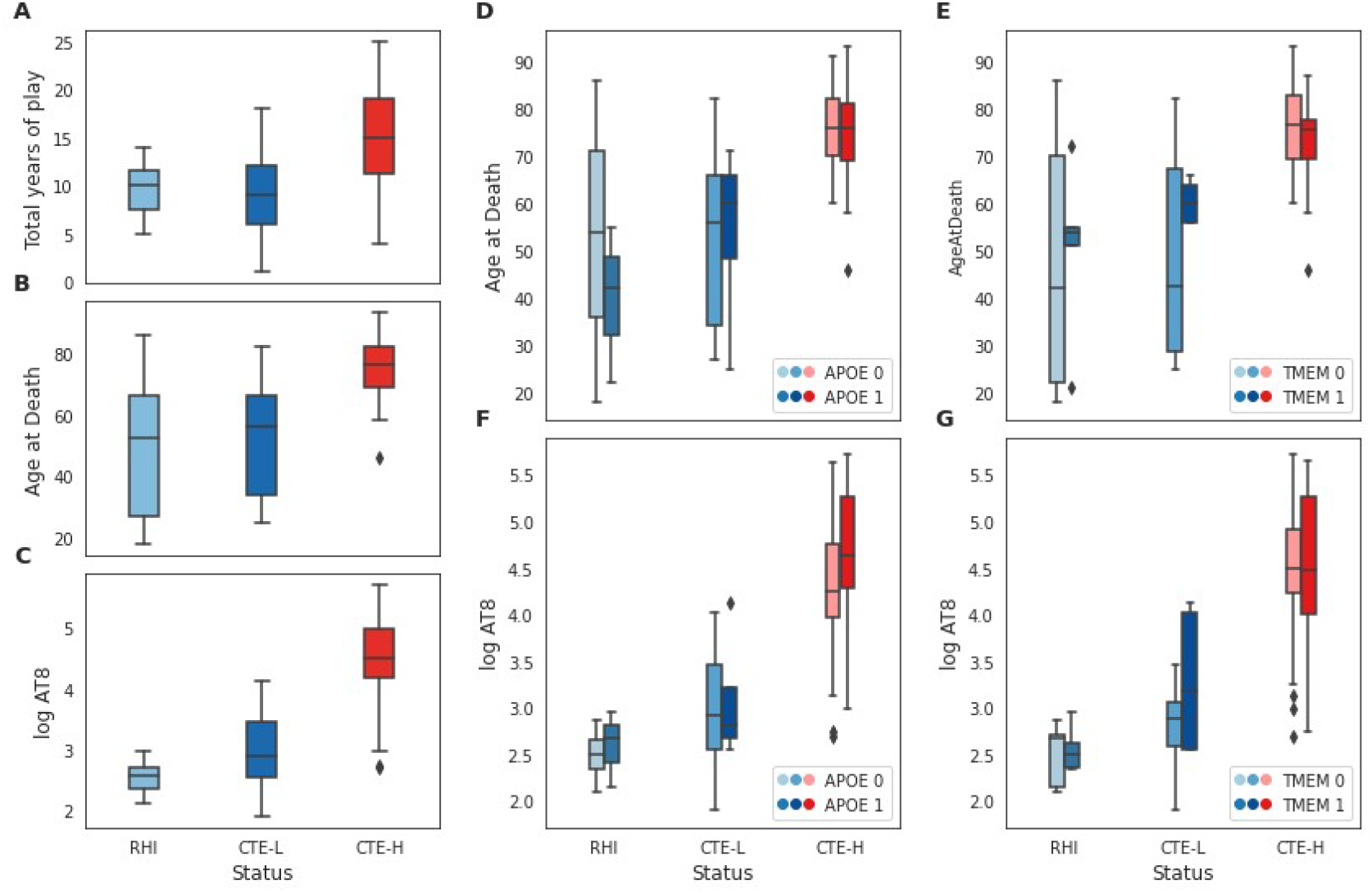
Sample characteristics for the variables examined in this study. A-C) Distribution of total years of play, age at death, and log AT8 histochemical quantification, respectively, for RHI, CTE-L, and CTE-H sample groups. D,E) Distribution of age at death for each sample group broken out into risk allele groups for APOE and TMEM106B, respectively. F,G) Distribution of log AT8 for each sample group as in D and E.

Five separate differential expression (DE) and subsequent gene set enrichment analyses (GSEA) using Gene Ontology (GO) annotations were conducted in this study. Each analysis sought to identify genes whose expression was associated with the different comparisons of interest in the same set of samples as described in Table 1. Specifically, separate DE models were conducted corresponding to case vs RHI control, number of years of play as a continuous variable, amount of tau pathology as measured by AT8 histochemistry, possession of one or more APOE e4 risk alleles, and possession of the TMEM106B risk allele (i.e. recessive model homozygous C for rs3173615). Samples were stratified into low (CTE-L, Stages I & II) and high (CTE-H, Stages III & IV) pathology groups and analyzed for each model separately. Table 1 contains sample count information and summary statistics for the six DE models reported in this study.

**Table 1.**
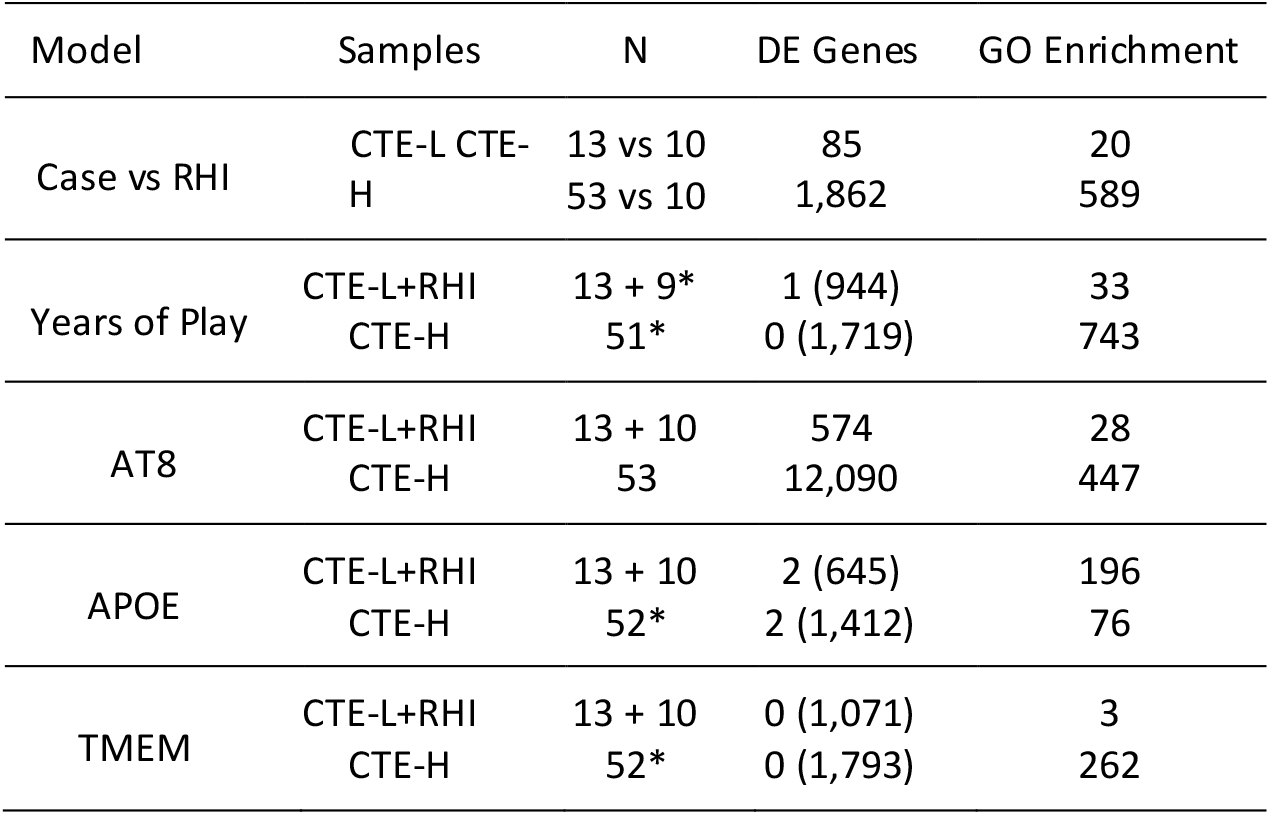
DE models and result statistics. The number of DE genes and enriched GO terms is reported for FDR < 0.1 (nominal p-value < 0.05 in parenthesis). All models include age at death, RIN, and sequencing batch as covariates in addition to the variable of interest. Case vs RHI is modeled as a binary variable with RHI samples as the reference group. Years of play and AT8 are modeled as continuous variables. *Some samples were omitted in certain comparisons due to outlier or missing values.

| Model | Samples | N | DE Genes | GO Enrichment |
| --- | --- | --- | --- | --- |
| Case vs RHI | CTE-L | 13 vs 10 | 85 | 20 |
|  | CTE-H | 53 vs 10 | 1,862 | 589 |
| Years of Play | CTE-L+RHI | 13 + 9* | 1 (944) | 33 |
|  | CTE-H | 51* | 0 (1,719) | 743 |
| AT8 | CTE-L+RHI | 13 + 10 | 574 | 28 |
|  | CTE-H | 53 | 12,090 | 447 |
| APOE | CTE-L+RHI | 13 + 10 | 2 (645) | 196 |
|  | CTE-H | 52* | 2 (1,412) | 76 |
| TMEM | CTE-L+RHI | 13 + 10 | 0 (1,071) | 3 |
|  | CTE-H | 52* | 0 (1,793) | 262 |

### 2.1 CTE versus RHI

Figure 2 depicts results from the case status DE models and subsequent gene set enrichment analysis. DE genes were primarily decreased in disease compared with RHI for both sample groups (Figure 2A,B). There was relatively consistent agreement in the direction of effect for genes when comparing the two analyses, and the common FDR significant DE genes were all changed in the same directions (marked genes in Figure 2C, listed in Table 2). However, there was little agreement in the direction of effect for gene set enrichment, where only four GO terms were significantly altered at FDR < 0.1 in both analyses, and three of those show an opposite direction of effect (Figure 2D and E). GO terms that trend toward significance (i.e. nominal p-value < 0.05) similarly showed substantial discordance in direction of effect for some GO terms (Figure 2E), where processes related to immune response, inflammation, blood brain barrier, extracellular matrix/membrane, and metabolism were increased in severe disease but decreased in mild disease. Processes increased in both were related to interaction with protein processing, metabolism, neuronal functions, and metal ion homeostasis, while those decreased in both involve mostly ribosomal processes and transcription/translation (Figure 2E).

**Table 2.** Common significant genes for CTE-L and CTE-H vs RHI from Figure 2C. Genes were significant at FDR < 0.1 in both comparisons. Mean columns contain base mean count estimates for each gene (i.e. baseMean reported by DESeq2) l2fc columns contain log2 fold change estimates.

| ENSG | Symbol | CTE-L |  |  | CTE-H |  |  |
| --- | --- | --- | --- | --- | --- | --- | --- |
|  |  | Mean | l2fc | FDR | Mean | l2fc | FDR |
| ENSG00000199805 | RNU1-134P | 15.59 | -1.65 | 0.10 | 18.08 | -1.44 | 0.03 |
| ENSG00000130856 | ZNF236 | 586.02 | -0.63 | 0.09 | 572.11 | -0.71 | 0.01 |
| ENSG00000276975 | HYDIN2 | 234.10 | -0.78 | 0.07 | 243.93 | -0.93 | 0.03 |
| ENSG00000286021 | AC112481.2 | 161.72 | -0.94 | 0.09 | 116.82 | -1.17 | 0.03 |
| ENSG00000244625 | MIATNB | 79.93 | -0.60 | 0.09 | 69.15 | -0.57 | 0.10 |
| ENSG00000233012 | HDAC1P2 | 20.30 | -1.36 | 0.07 | 21.60 | -1.15 | 0.02 |
| ENSG00000276002 | RF00017 | 8.09 | -2.38 | 0.04 | 11.42 | -1.32 | 0.09 |
| ENSG00000152208 | GRID2 | 154.23 | -1.10 | 0.04 | 134.44 | -1.35 | 0.00 |
| ENSG00000279557 | AC010435.1 | 17.62 | -1.52 | 0.04 | 21.08 | -0.98 | 0.07 |
| ENSG00000259706 | HSP90B2P | 71.62 | -1.19 | 0.09 | 82.97 | -1.28 | 0.05 |
| ENSG00000110975 | SYT10 | 55.42 | -1.28 | 0.09 | 47.40 | -1.75 | 0.00 |
| ENSG00000140992 | PDPK1 | 1789.15 | -0.54 | 0.09 | 1420.78 | -0.64 | 0.01 |
| ENSG00000265010 | AC087301.1 | 14.45 | -1.48 | 0.07 | 12.72 | -1.68 | 0.00 |
| ENSG00000273136 | NBPF26 | 398.68 | -1.36 | 0.04 | 493.63 | -1.20 | 0.07 |
| ENSG00000236829 | Z97634.1 | 120.46 | -1.06 | 0.09 | 125.60 | -0.89 | 0.06 |
| ENSG00000100218 | RSPH14 | 82.71 | -0.61 | 0.09 | 64.27 | -0.58 | 0.03 |
| ENSG00000196141 | SPATS2L | 1798.83 | -0.54 | 0.09 | 1558.49 | -0.50 | 0.02 |
| ENSG00000196290 | NIF3L1 | 266.39 | 0.67 | 0.09 | 168.84 | 0.68 | 0.06 |

**Figure 2.**
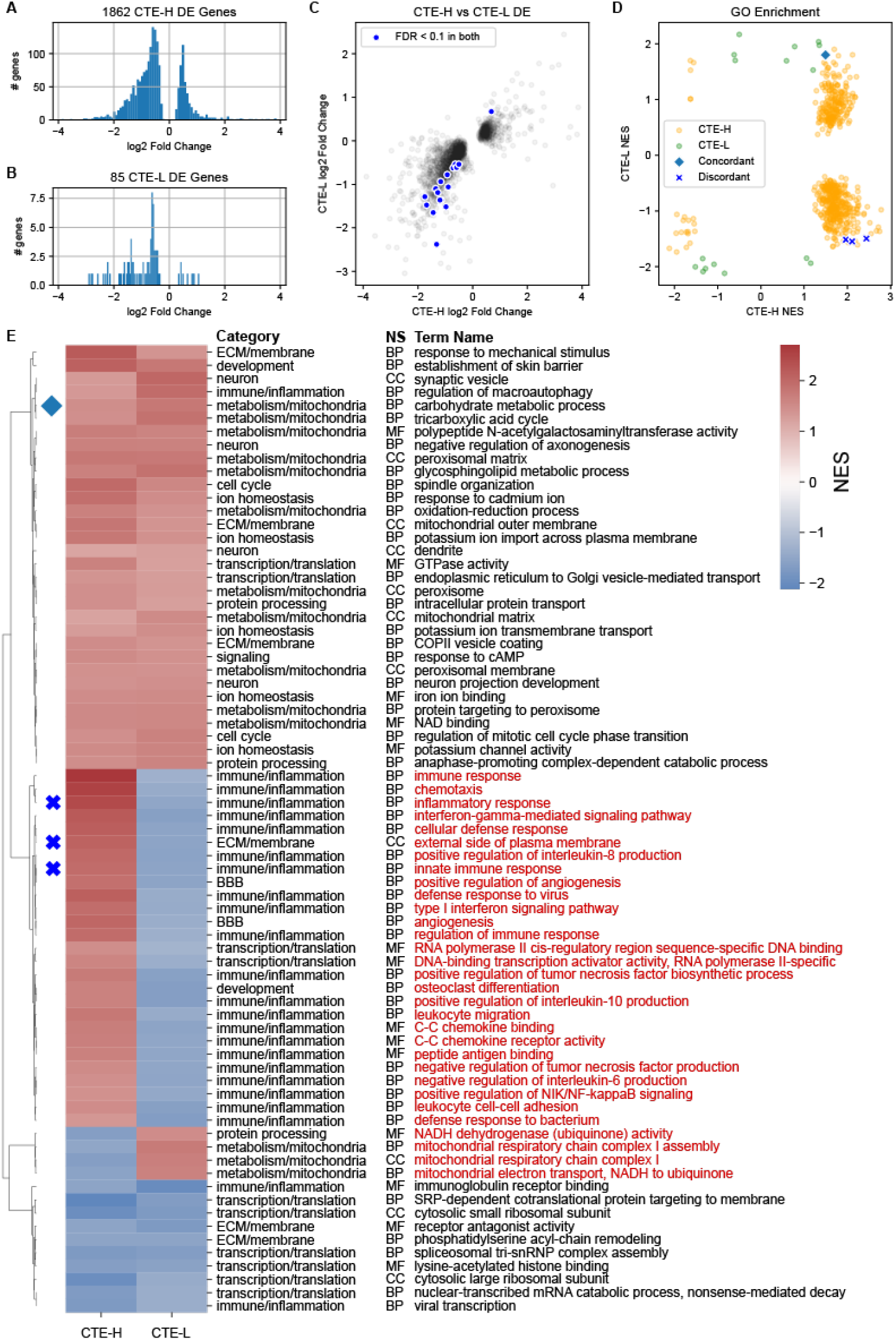
DE statistics for case versus RHI comparisons. Early and late stage CTE vs RHI controls showed general concordance in direction of DE genes, but mixed agreement on the biological process level. A-B) Distribution of log2 fold change for DE genes with FDR < 0.1 for both CTE-H vs RHI and CTE-L vs RHI, respectively. C) Log2 fold change values for DE genes with FDR < 0.1 in either CTE-H or CTE-L analyses. D) Normalized Enrichment Scores (NES) from Gene Set Enrichment Analysis (GSEA) of GO terms from the c5 MSigDB curated GO annotation at FDR < 0.1. Gene sets significant in both analyses are highlighted and colored based on concordance (i.e. same or different direction of effect). E) Hierarchically clustered heatmap of NES for enriched GO terms from D that have nominal p-value less than 0.05 in both CTE-H vs RHI and CTE-L vs RHI. Diamond and X markers correspond to GO terms significant at FDR < 0.1 in both from D. GO term names are colored red if the direction of effect is discordant between analyses. NS - GO namespace of corresponding term: BP biological process, CC - cellular component, MF - molecular function.

To understand which processes were unique to CTE-H or CTE-L, we filtered the GO terms to include only those with FDR < 0.05 in one analysis and nominal p-value > 0.25 in the other. This strategy identified 290 and 4 GO terms that were strongly enriched in CTE-H vs RHI and CTE-L vs RHI, respectively. The numbers of these uniquely enriched GO terms as well as the 41 with nominal p-value < 0.05 in both analyses depicted in Figure 2E organized by category were in Table 3. Immune and inflammation processes have the highest number of enriched GO terms and were represented in both analyses, with 15 terms in common between them. All biological categories were implicated by both analyses except for cytoskeleton, apoptosis, and signaling terms, which were only identified when comparing CTE-H and RHI.

**Table 3.** Counts of enriched GO terms grouped by high level category uniquely significant in CTE-L, in CTE-H, or implicated by both sample groups. ECM and BBB stand for extracellular matrix and blood brain barrier, respectively. ‘other’ contains GO terms not easily grouped into the other categories. GO term categories used were found in Supplemental Table S1.

| trend category | CTE-L | Both | CTE-H |
| --- | --- | --- | --- |
| cytoskeleton | 0 | 0 | 8 |
| apoptosis | 0 | 0 | 17 |
| signaling | 0 | 0 | 26 |
| neuron | 1 | 1 | 5 |
| cell cycle | 0 | 1 | 20 |
| development | 0 | 1 | 22 |
| protein processing | 0 | 1 | 45 |
| BBB | 0 | 2 | 4 |
| ECM/membrane | 0 | 2 | 32 |
| ion homeostasis | 0 | 3 | 5 |
| other | 0 | 3 | 6 |
| metabolism | 1 | 3 | 23 |
| transcription/translation | 0 | 9 | 16 |
| immune/inflammation | 2 | 15 | 76 |

To better summarize the large number of significantly enriched GO terms, we manually categorized GO terms found to be significant in any analysis into 13 high level categories, as depicted in Figure 3 (see Supplemental Material S1 for a complete categorization table). This categorization strategy enables concise comparison of different biological processes between groups, including direction of effect. The clustered heatmap of Normalized Enrichment Scores (NES) for GO terms significant at FDR *<* 0.1 in either CTE-H vs RHI or CTE-L vs RHI depicted in Figure 3A shows that there were terms which trend in both concordant and discordant directions of effect, and there is no obvious consistency in the concordance pattern from the perspective of categories. When the significant terms were grouped by category as in Figure 3B, immune/inflammatory processes appear as the most frequently increased in CTE-H vs RHI, while these processes appear decreased in CTE-L vs RHI.

Due to their relevance to this disease context, the immune/inflammation and neuron categories were further divided into subcategories based on their biological role (Figure 3C-D). From Figure 3C, increased neuronal development terms comprise most of the significant terms between CTE-H and RHI, while a small number of increased synaptic processes were implicated by CTE-L vs RHI. Increased innate immune processes were the most numerous in the severe CTE group but were closely followed by immune cell migration, cytokine-related inflammation, and apoptotic processes, while the few significant terms in CTE-L vs RHI suggest decreased phagocytosis, innate immune, antigen presentation, and adaptive immune processes (Figure 3D).

**Figure 3.**
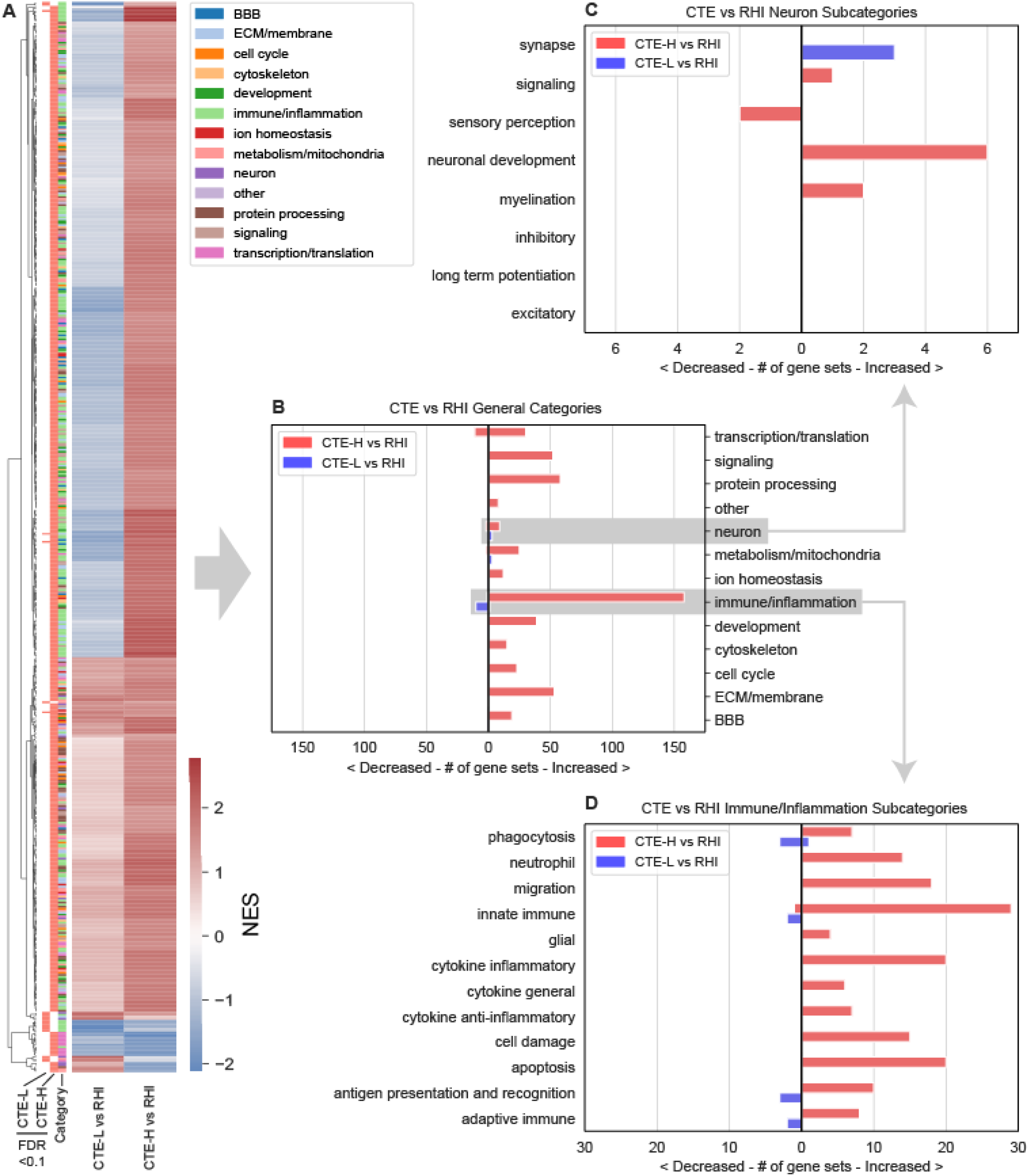
Enriched GO terms for case versus control. Detailed GO term enrichment of early and late CTE vs RHI showed concordant neuronal processes and opposite direction of effect for inflammatory processes. A) Heatmap of normalized enrichment scores (NES) for enriched GO terms. Row color bar represents significant pathways respective of columns (salmon colored bars to right of dendrogram indicate corresponding NES is FDR < 0.1) Rightmost color bar represents the GO category as listed in the legend. B) Number of significant GO terms from A grouped by category. Terms with positive, negative NES (red, blue in A respectively) are plotted as bars to the right and left of 0, respectively. C) Significant GO terms from the neuron category in B grouped by subcategory. D) Significant GO terms from the immune/inflammation category in B grouped by subcategory.

### 2.2 Association with Total Years of Play

We next sought to identify genes associated with exposure as measured by total years of play stratified into early (CTE-L+RHI) and late (CTE-H) sample groups. We chose to group CTE-L and RHI samples together in this analysis for three reasons. First, a goal of this study is to identify potential early changes in result of exposure to repetitive head trauma. Although CTE-L and RHI are pathologically distinct, they represent a similar level of exposure in years of play and age at death (Figure 1A,B), and therefore we hypothesize that similar genes are involved. Second, the brain tissue was taken from the gyral crest of the prefrontal cortex, which tends to be relatively spared of pathology in early disease compared with late, thereby avoiding most effects caused directly by pathology that likely influence the CTE-H samples. Last, we wanted to maximize our statistical power to detect differences with the total years of play variable by combining into a larger sample size.

Figure 4A and B compare DE genes and enriched GO terms for genes associated with total years of play, respectively. We note that there were no significant DE genes at FDR < 0.1 for either analysis (4A, Table 1), and unlike disease vs RHI DE genes, there is no apparent relationship between the direction of effect of these genes between CTE-L+RHI and CTE-H, and only 3 genes were nominally significant in both analyses (Figure 4A). GO term enrichment, on the other hand, identified many significant enriched terms at FDR < 0.1 that show consistently similar direction of effect of enriched terms, and all terms that are significant in both analyses are increased (Figure 4C, Table ??). Most of these common GO terms are related to immune response and inflammation, but unlike with comparison of disease vs RHI, all are increased in both CTE-L+RHI and CTE-H (Figure 4D). With the exception of neuronal processes in CTE-L+RHI, all implicated GO terms are positively associated with years of play in both sample groups.

**Figure 4.**
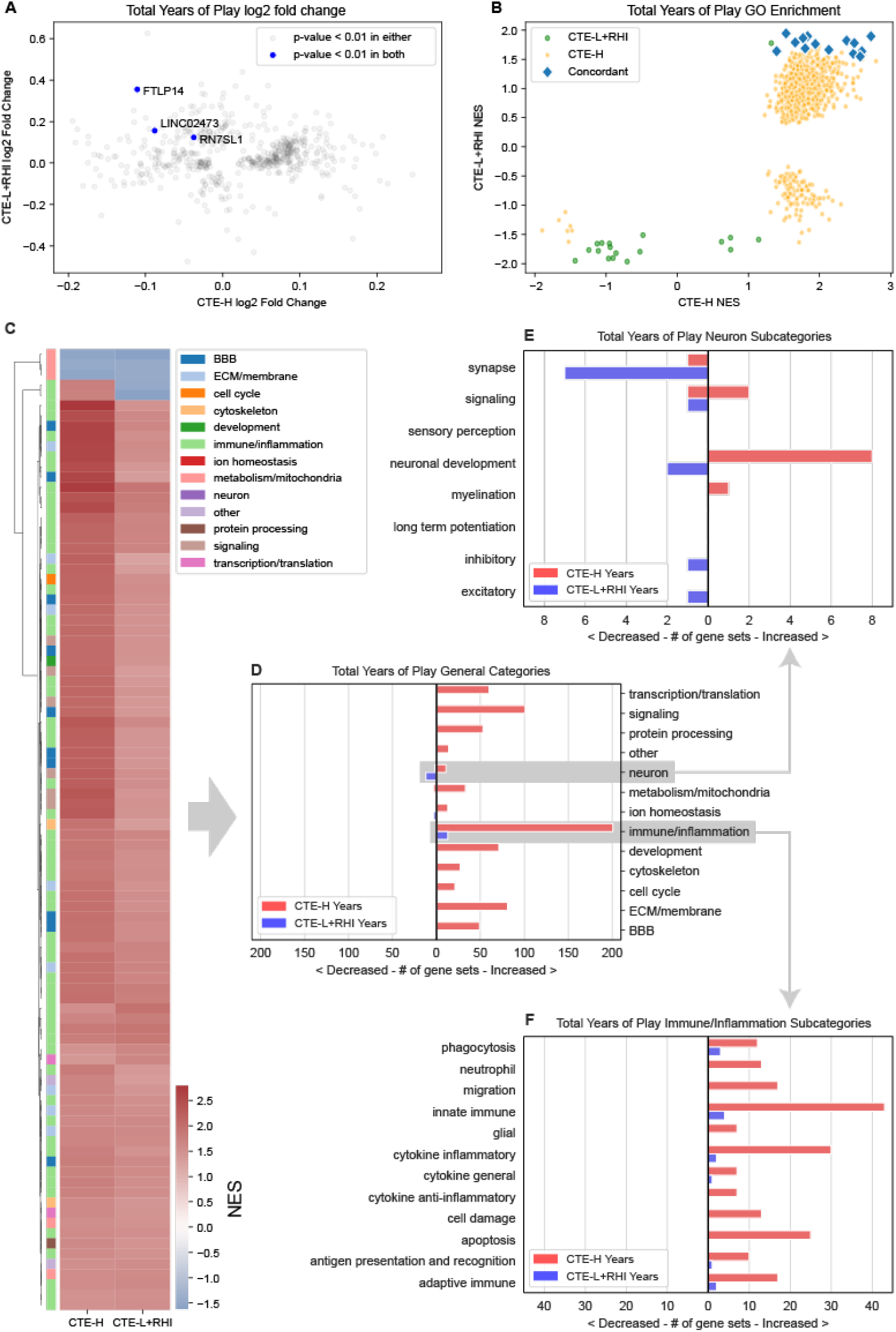
DE and GSEA statistics for genes associated with years of play. There were no FDR significant DE genes in either group but many significant gene sets in common that showed complete concordance in direction of effect between RHI+CTE-L and CTE-H. A) Log2 fold change estimates of genes associated with years of play for CTE-L+RHI (Low) and CTE-H (High) sample groups. As no genes were significant after adjusting for multiple hypotheses, genes with nominal p-value less than 0.01 are plotted. B) Gene set enrichment analysis of GO terms using log2 fold change. Gene sets are Concordant if they significant at FDR < 0.1 in both comparisons and are modified in the same direction. C) Clustered heatmap of enriched GO terms for both CTE-H and CTE-L+RHI associated with total years of play at nominal p-value < 0.05 in both. D) GO terms from C grouped by category. E) Neuron category terms. F) Immune/Inflammation category terms.

### 2.3 Association with Tau Pathology

We next sought to identify genes associated with tau protein aggregation as measured by immunohistological quantification of phospho-tau (AT8). AT8 immunostaining values were log transformed to attain normally distributed values. We again group CTE-L and RHI together for reasons described above. Although the amount of AT8 differs between CTE-L and RHI, the amount of tau pathology is more similar between CTE-L and RHI than CTE-H (Figure 1C, note log scale).

A substantial number of DE genes were associated with log AT8 at FDR < 0.1 for both CTE-L+RHI and CTE-H groups, and a large number of genes were significantly associated with both (Figure 5A). Strikingly, nearly all genes significant DE genes show an opposite direction of effect when comparing CTE-L+RHI and CTE-H. This inverse relationship is also observed in the enriched GO terms induced by the DE genes (Figure 5B), where all common enriched terms are discordant in direction of effect. The inverse relationship is also visible in Figure 5C, where immune/inflammation processes are primarily up in CTE-H but trend down in CTE-L+RHI, and the reverse is true of many other terms. This is shown clearly in Figure 5D, where many immune/inflammation terms are increased in association with AT8 in the CTE-H group, similar to the association seen with total years of play. However, CTE-H also shows an equally large number of neuronal terms that are decreased with increasing amounts of tau and, curiously, these processes appear to be positively associated with AT8 in the CTE-L+RHI group.

**Figure 5.**
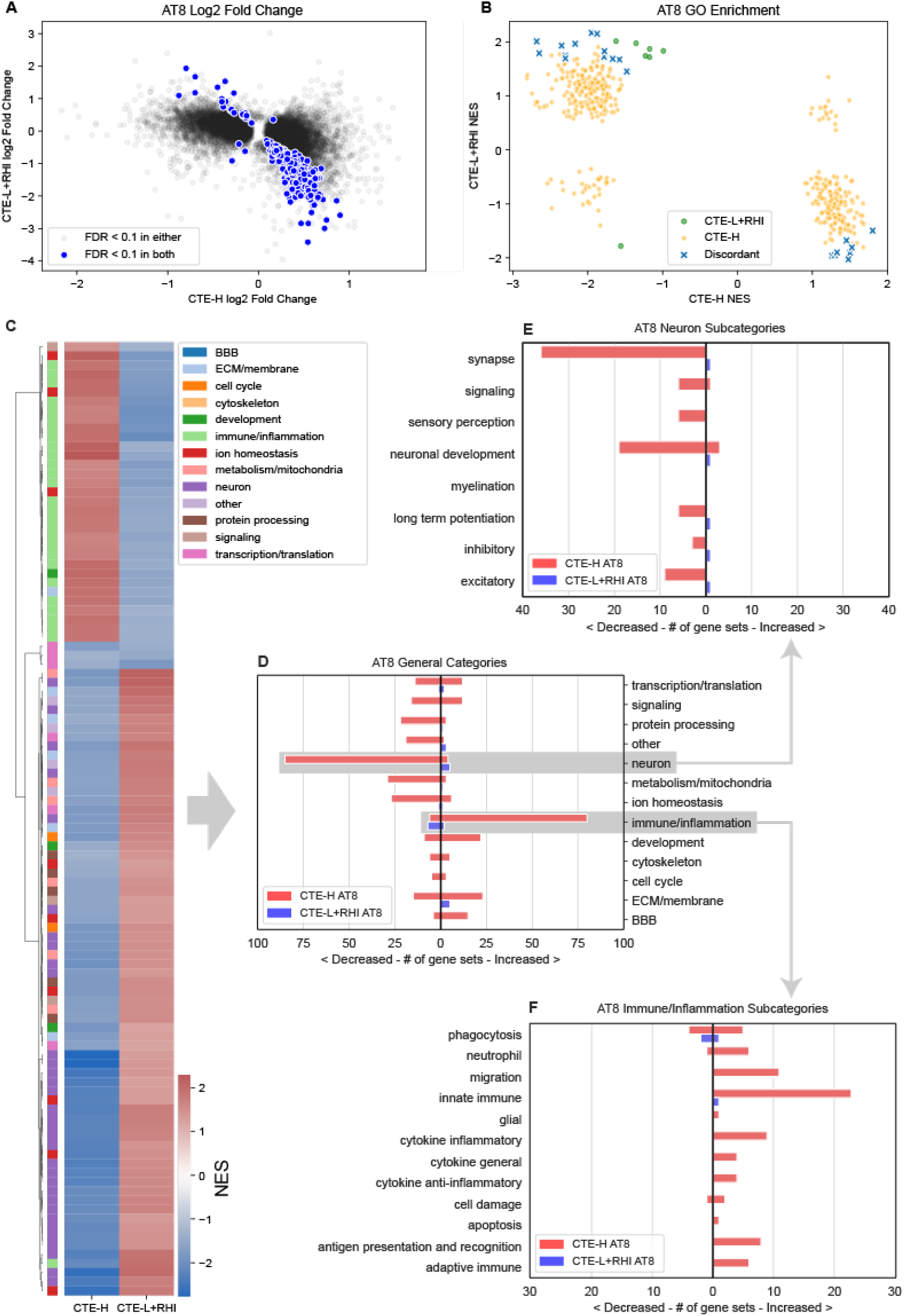
DE and GSEA statistics for genes associated with log AT8. There were many FDR significant DE genes and gene sets associated with AT8 that showed almost complete discordance in direction of effect between RHI+CTE-L and CTE-H. A) Log2 fold change estimates of genes associated with AT8 for CTE-L+RHI (Low) and CTE-H (High) sample groups. B) Gene set enrichment analysis of GO terms using log2 fold change. Gene sets were Discordant if they were significant in both comparisons and are modified in opposite directions. C) Clustered heatmap of enriched GO terms for both CTE-H and CTE-L+RHI associated with total years of play at nominal p-value < 0.05 in both. D) GO terms from C grouped by category. E) Neuron category terms. F) Immune/Inflammation category terms.

### 2.4 Risk Variant Effects

We next investigated how risk variants of APOE and TMEM106B affect early and late CTE disease stages. A dominant encoding for the APOE e4 allele was used to separate samples in each sample group, i.e. individuals with at least one e4 allele were included in the risk group. The TMEM106B SNP rs3173615 G allele has been shown to be protective^7^. Thus, individuals homozygous for the non-protective C allele were grouped to form the risk carriers (i.e. a recessive encoding for the non-protective allele). This encoding makes comparing the results of the two variants more straightforward, as association of genes and GO terms have the same interpretation with respect to the risk carriers.

DE analyses were conducted comparing the risk vs non-risk subjects within each of the CTE-L+RHI and CTE-H groups, resulting in four sets of DE results. Very few genes were significantly associated at FDR *<* 0.1, where only 2 met this significance level in the APOE comparisons and none for TMEM106B (see Table 1). However, 196 and 76 GO terms were significantly associated at FDR *<* 0.1 for CTE-L+RHI and CTE-H, respectively, with APOE risk, and 3 and 262 were significantly associated with these in TMEM106B risk. It is interesting to note that the number of significant GO terms is higher for APOE risk in the CTE-L+RHI group, which has the smallest number of samples, and also higher in the CTE-H group for TMEM106B risk, where the remaining two analyses yielded relatively few results.

We next compared the log2 fold change associated with the risk allele group of genes between variants and sample groups (Figure 6A-D). In principle, if the risk variants modulate disease risk using a common mechanism, the DE genes identified for the different risk genes and sample groups should show some similarity. Very few nominally significant genes overlapped between any of the comparisons, as illustrated in the Venn diagrams of Figure 6. However, similar to the inverse relationship observed between CTE-L+RHI and CTE-H in the AT8 comparison, we observed that the effect size of the overlapping genes was inverted between these groups for APOE risk carriers (Figure 6A scatter plot). The relationship was less consistent when comparing TMEM106B risk carriers between CTE-L+RHI and CTE-H, where there were both concordant and discordant directions of effect in the common genes (Figure 6B scatter plot). Curiously, an inverse relationship is also observed when comparing the APOE and TMEM106B risk carriers within the CTE-L+RHI group (Figure 6C scatter plot). There is no consistent relationship between the genes for the two risk carriers within CTE-H (Figure 6D scatter plot).

**Figure 6.**
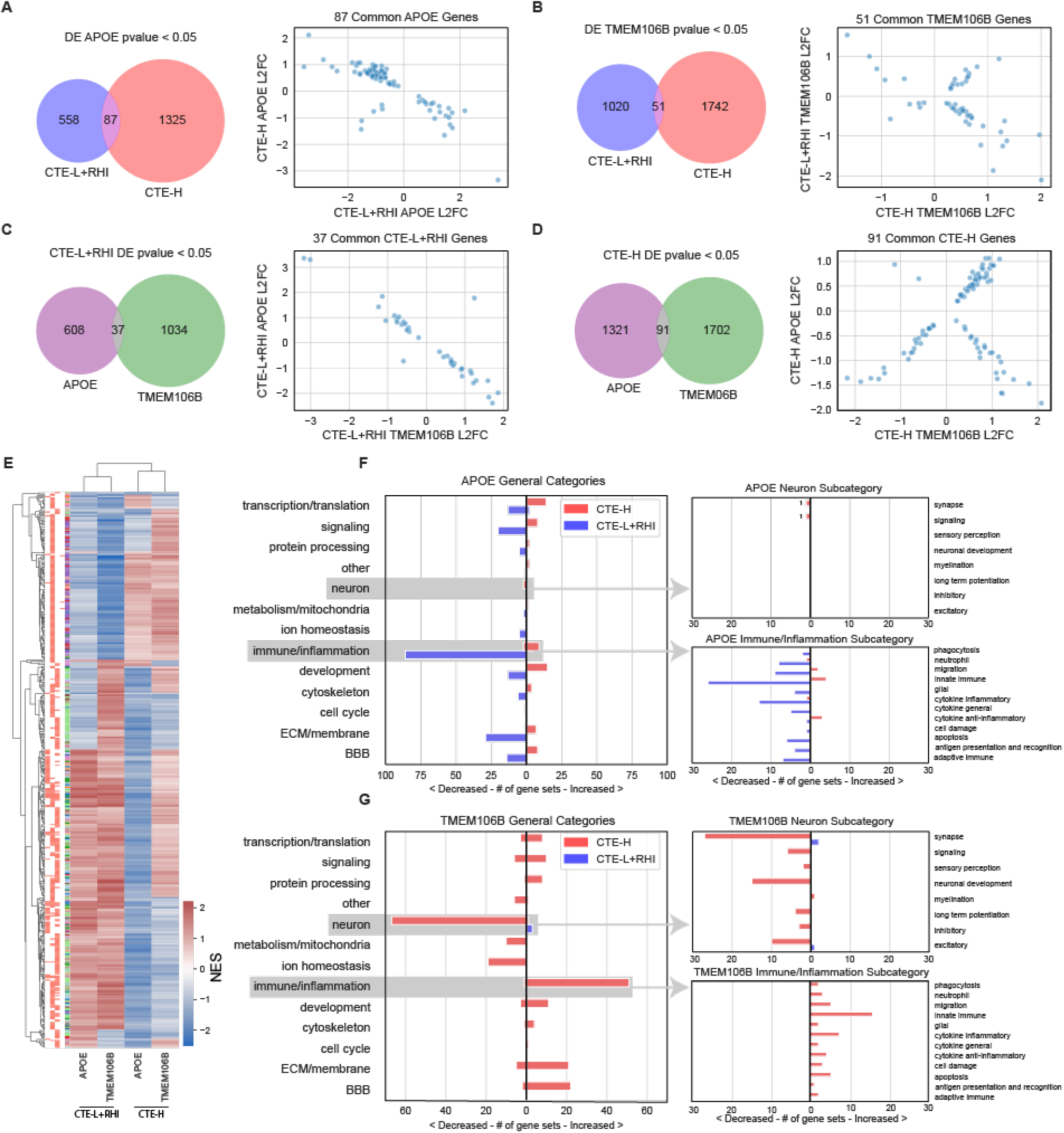
DE and GSEA statistics for genes and gene sets associated with APOE *ε*4 and TMEM106B risk alleles. Genes associated with risk alleles in both sample groups were largely disjoint, and gene sets associated with APOE and TMEM106B risk alleles were primarily found in CTE-L+RHI and CTE-H, respectively. A-B) Venn diagram of nominally significant (p-value < 0.05) gene overlap and scatterplot of L2FC of CTE-L+RHI vs CTE-H sample groups for APOE (A) and TMEM106B risk allele (B), respectively. C-D) Venn diagram of nominally significant (p-value < 0.05) gene overlap and scatterplot of L2FC of APOE *ε*4 vs TMEM106B risk allele for CTE-L+RHI (C) and CTE-H (D), respectively. E) NES heatmap of GO terms significant at FDR < 0.1 in at least one condition. Bars on left of the heatmap indicate significance of corresponding NES in the heatmap columns respectively, GO IDs that were significant at FDR < 0.1 is in red. F-G) GO pathways that were FDR < 0.1 grouped by category. Values left of zero corresponds to number of significant pathways with negative NES. Values right of zero corresponds to number of significant pathways with positive NES.

Similar to the gene set enrichment analyses presented earlier, there was a mix of concordance and discordance in the direction of effect of GO terms implicated by the DE genes. The clustered heatmap in Figure 6E depicting Normalized Enrichment Scores for each analysis showed little consistent pattern of concordance between them, except to note that the sample groups clustered together more closely than the variant groups. The discordance in direction of effect for APOE variants across sample groups was also apparent (1st and 3rd column of heatmap), consistent with the inverted gene expression relationship from Figure 6A. More generally, the patterns observed in comparing concordance between pairs of columns in the heatmap are consistent with the scatter plots by inspection.

There was a surprising number of significantly enriched GO terms associated with APOE risk carriers in the CTE-L+RHI group in Figure 6F. No other analysis conducted by this study found so many significant results in this sample group, which was notable considering this was the sample group with the smallest sample size. Many GO terms in various categories, most notably neuron, are negatively associated with APOE risk carriers, meaning neuronal gene expression overall was decreased in early individuals who have the APOE risk allele. In contrast to the results from APOE, the pattern of biological processes implicated by TMEM106B were more consistent with other comparisons made in this study, namely that CTE-H exhibited the majority of the significantly associated GO terms which were primarily composed of decreased neuron and increased immune/inflammation categories (Figure 6G).

### 2.5 Validation of Associations with qPCR

To provide orthogonal validation of these findings, we examined previously generated qPCR expression quantification on an a priori set of 33 genes known to be implicated in CTE (full gene list in Supplemental Table S2) in a subset of 54 participants (37 CTE-H, 9 CTE-L, and 8 RHI). These genes were chosen based on our previous investigations as being involved more generally in neurodegenerative disease processes and as such many were not significantly DE in our mRNASeq CTE analyses (see Supplemental Table S3). We therefore focused on comparing the direction of effect (i.e. log2 fold change) between the qPCR and DE genes to assess concordance and tabulated the results into confusion matrices found in Table 5.

**Table 4.** DE models and result statistics. The number of DE genes and enriched GO terms is reported for FDR < 0.1. Number of DE genes at nominal pvalue < 0.05 are parenthesized. Model samples were grouped by their risk variant category. All models include age at death, RIN, and sequencing batch as covariates in addition to the variable of interest. AT8 was modeled as continuous variables.

| Model | Samples | N | DE Genes | GO Enrichment |
| --- | --- | --- | --- | --- |
| APOE 0 | CTE-L+CTE-H | 9 + 29 | 5,866 | 385 |
|  | CTE-L | 9 | 19 | 77 |
|  | CTE-H | 29 | 3,401 | 283 |
| APOE 1 | CTE-L+CTE-H | 4 + 23 | 2,649 | 409 |
|  | CTE-H | 23 | 503 | 321 |
| TMEM 0 | CTE-L+CTE-H | 5 + 18 | 7,919 | 490 |
|  | CTE-L | 5 | 1 (116) | 25 |
|  | CTE-H | 18 | 7,159 | 500 |
| TMEM 1 | CTE-L+CTE-H | 8 + 34 | 3,117 | 292 |
|  | CTE-H | 34 | 449 | 128 |

**Table 5.**
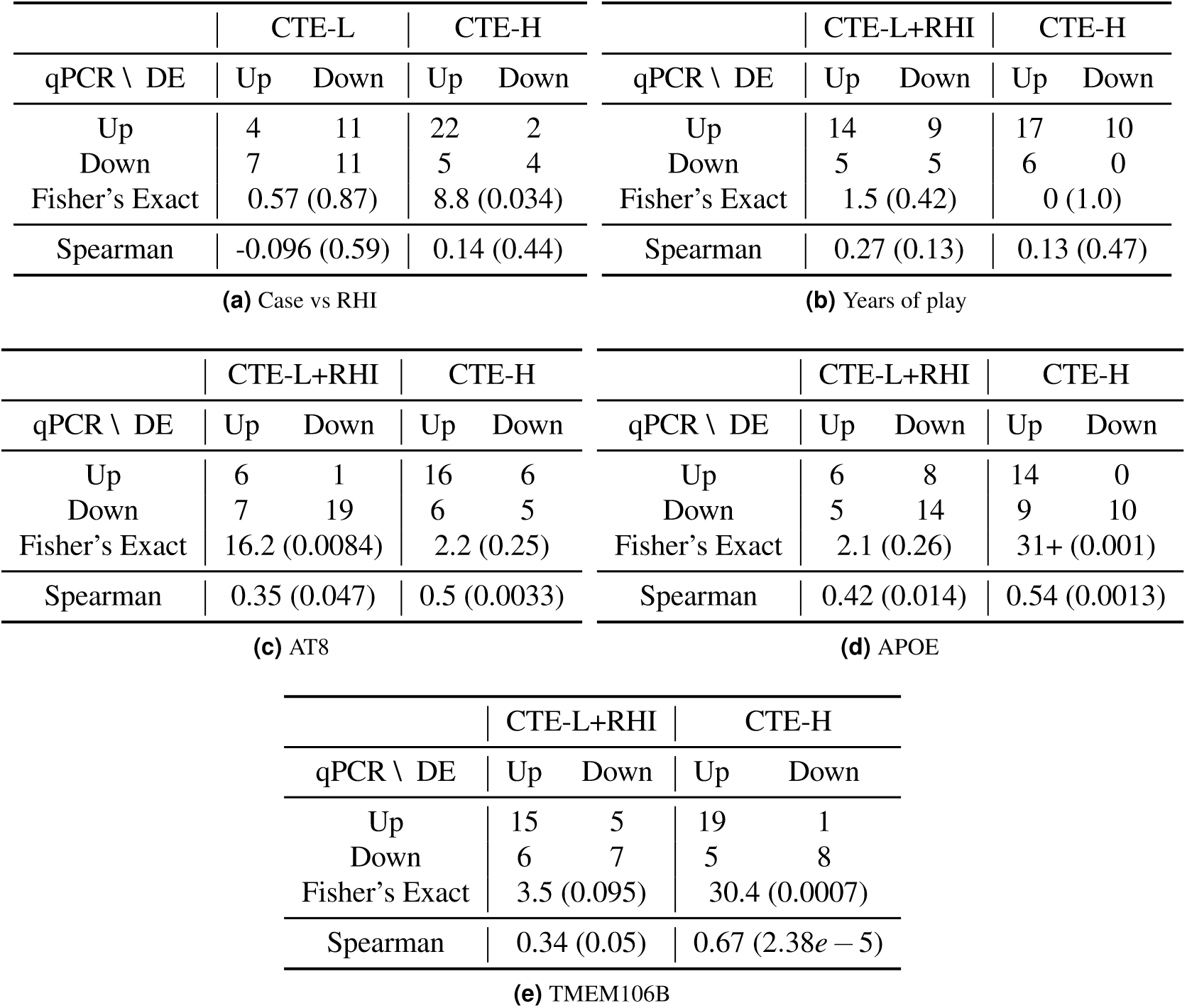
DE genes vs qPCR log2 fold changes for all five models reported in this study. Top two rows (qPCR) and columns (mRNASeq) of each table show directional overlap (i.e. positive vs negative log2 fold changes) of 33 genes where both mRNASeq counts and qPCR data are available. Fisher’s Exact rows report odds ratio and associated right-tailed p-value for the contingency tables above. Spearman rows report Spearman correlation and associated p-value for comparing the log2 fold changes between mRNASeq and qPCR.

|  | CTE-L |  | CTE-H |  |
| --- | --- | --- | --- | --- |
| qPCR \ DE | Up | Down | Up | Down |
| Up | 4 | 11 | 22 | 2 |
| Down | 7 | 11 | 5 | 4 |
| Fisher's Exact | 0.57 (0.87) |  | 8.8 (0.034) |  |
| Spearman | -0.096 (0.59) |  | 0.14 (0.44) |  |
| (a) Case vs RHI |  |  |  |  |

**Table 5.** DE genes vs qPCR log2 fold changes for all five models reported in this study.
|  | CTE-L+RHI |  | CTE-H |  |
| --- | --- | --- | --- | --- |
| qPCR \ DE | Up | Down | Up | Down |
| Up | 6 | 1 | 16 | 6 |
| Down | 7 | 19 | 6 | 5 |
| Fisher's Exact | 16.2 (0.0084) |  | 2.2 (0.25) |  |
| Spearman | 0.35 (0.047) |  | 0.5 (0.0033) |  |
| (c) AT8 |  |  |  |  |
| qPCR \ DE | Up | Down | Up | Down |
| Up | 15 | 5 | 19 | 1 |
| Down | 6 | 7 | 5 | 8 |
| Fisher's Exact | 3.5 (0.095) |  | 30.4 (0.0007) |  |
| Spearman | 0.34 (0.05) |  | 0.67 (2.38e−5) |  |
| (e) TMEM106B |  |  |  |  |
| qPCR \ DE | Up | Down | Up | Down |
| Up | 14 | 9 | 17 | 10 |
| Down | 5 | 5 | 6 | 0 |
| Fisher's Exact | 1.5 (0.42) |  | 0 (1.0) |  |
| Spearman | 0.27 (0.13) |  | 0.13 (0.47) |  |
| (b) Years of play |  |  |  |  |
| qPCR \ DE | Up | Down | Up | Down |
| Up | 6 | 8 | 14 | 0 |
| Down | 5 | 14 | 9 | 10 |
| Fisher's Exact | 2.1 (0.26) |  | 31+ (0.001) |  |
| Spearman | 0.42 (0.014) |  | 0.54 (0.0013) |  |
| (d) APOE |  |  |  |  |

The level of agreement across sample groups and models between qPCR and mRNASeq DE varied. Some comparisons showed very little agreement in the direction of effect, particularly the CTE-L vs RHI and total years of play models. Note from the mRNASeq analysis of total years of play above (Figure 4) that there were very few DE genes implicated and very little agreement on the direction of effect, which was consistent with the results of Table 5b). On the other hand, some comparisons show very high concordance, the most noteworthy being those for AT8 (Table 5c). Here the unexpected inverse relationship of CTE-L+RHI and CTE-H with AT8 (see Figure 5A) was also observed; note most genes are decreased and increased in CTE-L+RHI and CTE-H, respectively, and the Fisher’s Exact test for CTE-L+RHI attained significance and CTE-H trends toward significance. The Spearman correlation of log2 fold changes for AT8 are modest but significant at p-value < 0.05. The TMEM106B risk allele comparisons (Table 5e) also show substantial agreement. Overall, the concordance of results from qPCR and mRNASeq was remarkable, especially considering the genes were chosen independently.

### 2.6 Comparative Analysis

To aid in summarizing results presented in this study, subpanels of GO category enrichment for each of the five primary analyses were included in Figure 7. These plots depict the number of significantly enriched GO terms grouped by high level category as in each analysis specific figures presented earlier. The subfigures have been annotated to emphasize several salient patterns observed across analyses. Increased immune/inflammation processes were implicated in all comparisons involving CTE-H except with APOE risk. CTE vs RHI and total years of play analyses (Figure7A,C) were nearly absent of neuronal processes, while in AT8 and TMEM106B risk comparisons, neuronal processes were substantially decreased for CTE-H (Figure7B,D). Comparisons with CTE-L had very few enriched GO categories except for APOE risk, where there was a substantial decrease in immune/inflammatory categories.

**Figure 7.**
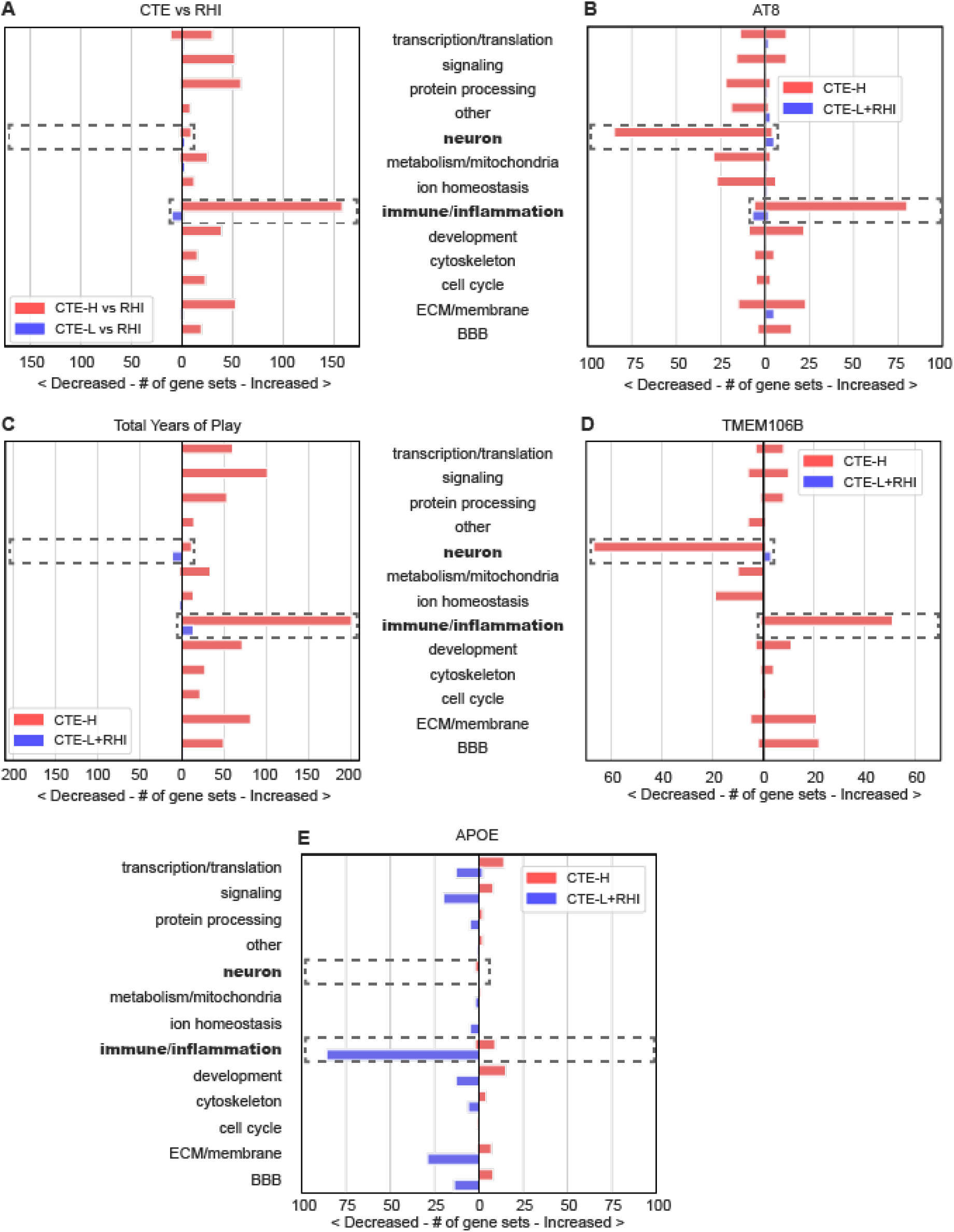
Enriched GO term categories for all five primary analyses presented in this work as found in previous figures. Increased inflammation was associated with all CTE-H analyses except APOE risk carriers, and decreased neuronal processes were only associated with AT8 and TMEM106B risk. A) CTE vs RHI, B) AT8, C) total years of play, D) TMEM106B risk carriers vs non-risk carriers, and E) APOE risk carriers vs non-risk carriers. Dashed boxes and bolded text are annotated to aid in interpretation.

## 3 Discussion

This study presents the largest genome-wide gene expression analysis of post-mortem human brain in CTE to date. We set out to identify gene expression patterns observed in early vs late disease as they relate to presence of pathology (i.e. case vs RHI), the amount of repetitive head impacts exposure (i.e. total years of play), quantitative measures of tau pathology in affected brain tissue, and genetic variants known to influence CTE symptoms and severity. We showed that there were substantial gene expression effects in individuals with severe CTE across all of these axes while the comparisons involving RHI and CTE-L had fewer significant results. Case vs control comparisons yielded mixed concordance in the direction of effect of implicated pathways, while processes associated with total years of play were in good agreement between early compared with late sample groups. In contrast, the processes associated with the amount of tau pathology had almost exactly opposite directions of effect between these groups. Furthermore, the DE genes associated with APOE and TMEM106B risk variants did not have a high degree of overlap.

In every comparison except APOE risk carriers, substantial neuroimmune and neuroinflammatory processes were positively associated with conditions that increase risk of severe disease in CTE-H subjects. Processes in the innate immune response subcategory were the most numerous, but many different components of the immune response, including cytokine activity and apoptosis, were also well represented in these comparisons. While some inflammatory processes implicated when comparing CTE-H APOE risk carrier groups were consistent with these findings with disease, RHI, tau pathology, and TMEM106B risk, the small number of processes identified may suggest that the effects of APOE risk variants largely precede the development of severe disease. This idea is further supported by the finding that the comparison of APOE risk carriers in CTE-L+RHI samples produced many significantly enriched gene sets, whereas no other comparison implicated nearly as many for this sample group. Curiously, these processes, most notably immune/inflammation, appear reduced in APOE risk carriers relative to non-risk carriers in this sample group, forming an almost exact mirror image of the increased processes observed in CTE-H. This suggests the possibility that the APOE risk variant may impair these processes in early disease, which in turn might predispose risk carriers to developing more severe pathology over time than non-risk carriers. An alternative explanation might be that the APOE risk allele causes an aberrant increase in the inflammatory state of the brain which is then compensated by homeostatic mechanisms that are competent in early disease but become less effective over time, leading to dramatically increased inflammation in late stage disease.

A second noteworthy pattern we observed is that neuronal gene expression decreases with increasing tau pathology and with the presence of TMEM106B risk. It is interesting to note that comparatively few neuronal processes are implicated when comparing CTE-H and CTE-L to RHI or when examining gene expression associated with total years of play, and almost none were associated with APOE risk status. Synapse was the neuronal subcategory with the largest number of enriched gene sets across these comparisons, followed by neuronal development. With the exception of CTE-L vs RHI, synaptic genes are decreased with increasing disease risk factors, suggesting that synaptic density is reduced in affected tissues, as has been recently shown in mouse models of repetitive head impact^15^. TMEM106B risk, in particular, was associated with decreased expression of many neuronal process pathways in CTE-H, which is consistent with previous reports demonstrating an association with decreased neuronal density ^30^. In addition, the previously demonstrated increased variation in synaptic density in CTE suggests that synapse turnover may be a feature of RHI and CTE and may be associated with greater variation in synapse related gene expression^16^.

The amount of head injury exposure as measured by total years of play appears to have only a weak effect on individual gene expression across individuals, but the biological processes they implicate are relatively consistent when comparing exposure groups. A given individual’s exposure to repetitive head impacts may have occurred many years prior to death when these samples were collected, and while we adjusted for the effects of age at death as well as possible, this duration paired with diverse life experiences and disease progression could easily distort any consistent signal on the gene level. The consistency between sample groups on the biological process level is therefore noteworthy and suggests there may indeed be a common response to exposure detectable even many years after exposure.

In contrast with years of play, tau pathology in the post-mortem samples captures an immediate condition of the brain when gene expression is measured. Indeed, the presence of tau pathology appears to have a strong effect on gene expression in both CTE-L+RHI and CTE-H sample groups. Thousands of genes and hundreds of enriched GO term gene sets are associated with AT8 immunopositivity, primarily implicating inflammatory and neuronal processes. The striking and nearly complete inverse relationship between both gene expression fold changes and gene set enrichment direction suggests a qualitatively different effect of pathology, or precursors to development of pathology, in individuals with low compared with high levels of exposure. Since the amount of tau pathology is relatively low but increased in CTE-L compared with RHI, this suggests a fundamentally different process affects these groups than in severe disease. With the exception of inflammation, development, and ECM/membrane, all biological processes trend as negatively associated with increasing amounts of tau. We recently found similar gene expression changes comparing sulcal vs gyral crest in dorsolateral prefrontal cortex in CTE subjects^17^.

The concordant effects of years of play and the discordant effects of tau explain the mixed concordance observed when comparing case vs RHI. Because the amount of tau pathology and years of play vary among individuals in both low and high exposure groups, we may therefore interpret the case vs RHI comparison as a convolution of these disparate effects. Although years of play and pathology are highly correlated, this motivates considering these as separate effects that modulate disease expression, which may have important therapeutic implications.

The relative lack of strong gene expression signal in APOE risk carriers is somewhat surprising, considering evidence of the role this gene is thought to play in the development of tau pathology in CTE and AD. Also surprising is the lack of overlap of DE genes and the discordance of the direction of effect between APOE and TMEM106B risk carriers vs non-risk carriers. Risk alleles of both genes increase the risk of developing severe CTE pathology, but the low overlap of the DE genes and biological processes suggests that the molecular mechanisms underlying this increased risk are distinct. As in the AT8 comparison, there is an inverse relationship between the nominally significant DE genes comparing CTE-L+RHI vs CTE-H APOE risk groups as well as TMEM106B vs APOE in the CTE-L+RHI sample group. The similarity of expression profiles between tau and TMEM106B does not appear to be driven by the amount of tau pathology, as the distribution of tau in the TMEM106B risk carriers and non-carriers does not significantly differ. The reasons for this are unclear, but further suggest a different molecular process is at play when considering the effects of these genes in combination with tau and exposure.

The relatively small number of FDR significant DE and GSEA results for the CTE-L vs RHI and CTE-L+RHI analyses in the study is likely due in part to low statistical power on account of relatively small sample size, but, since the results are sparse, the influence of false positives is minimal. On the other hand, the relative paucity of results compared with severe disease comparisons may indicate that early molecular changes in the brain of those with CTE are in fact similar to those who experienced a similar level of repetitive trauma exposure but lack specific CTE pathology. This would suggest a model where the pathology itself is a driver of molecular changes in later disease stages, which is supported by the observation that gene expression in many genes is associated with the amount of tau pathology.

The study has several important limitations. First, although we considered all viable CTE-L and RHI subjects available at the time of study, the number of these samples is relatively low for transcriptome wide study. While we attempted to mitigate this limitation by combining these two groups together when possible, further work is needed on a larger dataset to confirm the findings.

A second limitation is inherent to the nature of subjects affected by CTE and the unavoidable correlation of their characteristics. It is important to consider the inherent correlation of key characteristics of individuals affected by CTE, and relationships between the variables examined in this study and their manifestation in disease. There is substantial correlation between the severity of disease, the number of years of play, age at death, and amount of pathology present in the brain in CTE. This collinearity is further compounded by the presence of the APOE and TMEM106B risk alleles that modify these factors. CTE stage is determined in large part by the amount and localization of tau pathology, which increases with exposure, age, and risk variant status. It is also difficult to avoid confounding during sample selection, as lower stage disease will almost always correspond to some combination of these factors. Thus, it is difficult if not impossible to completely avoid this confounding and isolate the individual effects of these variables, even with much larger sample sizes than are presented here. It is therefore important to interpret the results of all of these variables together with this intrinsic confounding in mind and much larger sample sizes will be required to confidently disentangle the contribution of these factors. Generating expression data in a larger set of all three sample types will help clarify these ambiguities and is in progress.

In conclusion, these data present compelling evidence of widespread gene expression changes in late stage CTE and are less pronounced changes in early disease and those with repetitive head impacts exposure but without CTE pathology. Furthermore, these results suggest individuals with low exposure and little to no pathology experience a different set of molecular processes than those with late disease as well as a distinct association with tau pathology and genetic risk factors. Therapeutics and biomarkers developed as targets with late-stage signatures might not be effective for individuals early in the progression of disease. Future studies should endeavor to further characterize the active disease process in younger individuals with less exposure.

## 4 Methods

### 4.1 Sample Characteristics

All brains were collected and maintained by the Boston University Alzheimer’s Disease & CTE Center Brain Bank. This study has been approved by the VA Bedford Healthcare System and Boston University School of Medicine Institutional Review Boards. Participants with a history of contact sports participation were drawn from the Veterans Affairs-Boston University-Concussion Legacy Foundation (VA-BU-CLF) brain bank^18^. Participants were excluded if they lacked fresh frozen prefrontal cortex, died from drug overdose, hanging, or gunshot wound to the head, or had motor neuron disease or other neurodegenerative disease in the absence of CTE. Complete sample statistics are included in Table 6.

**Table 6.**
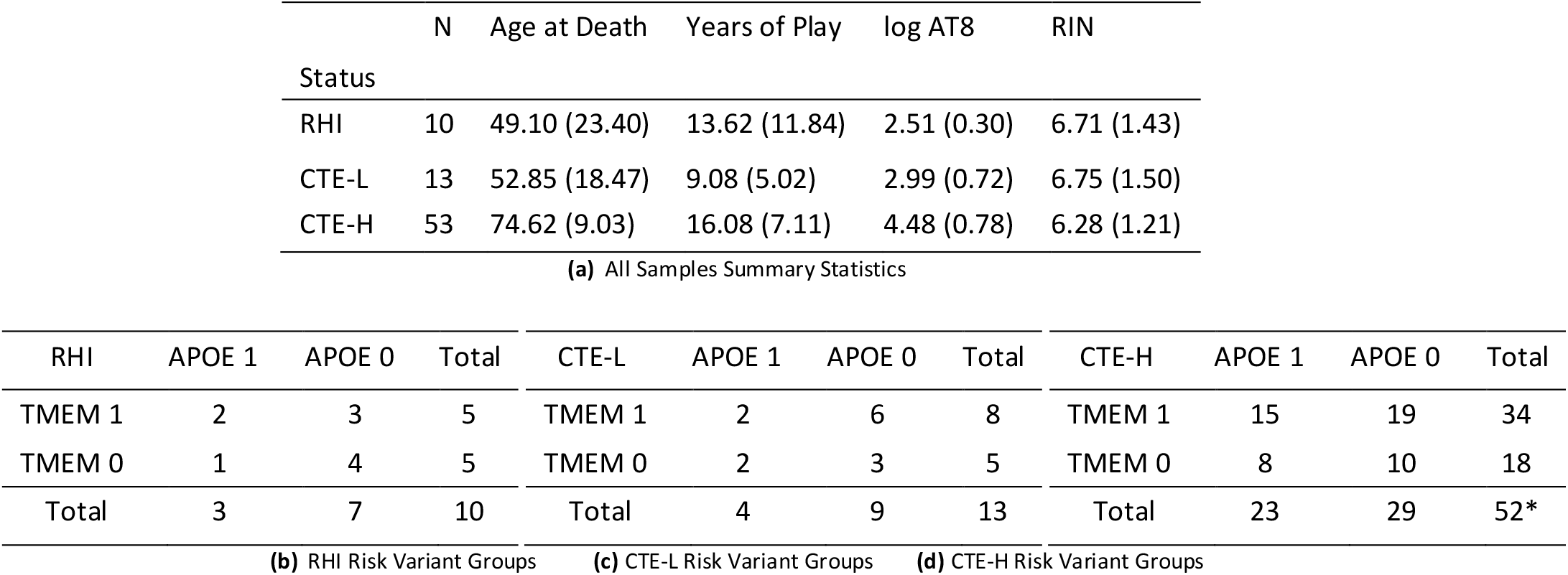
Sample statistics. Primary and parenthesized figures are mean and standard deviation for the given covariate, respectively. AT8 is measured in number of chromagenic pixels from histological scan using the AT8 antibody and are log transformed for use in this study. RNA Integrity Number (RIN) of the extracted RNA sample from each brain was assessed by an Agilent Bioanalyzer instrument. A) All sample statistics. Numbers in parentheses are standard deviations. B-D) Sample statistics for each group categorized by risk variants. Athletes with no APOE4 allele were classified as APOE 0 and those with at least one APOE4 allele was classified as APOE 1. Additionally, athletes with a G in TMEM106B rs3173615 were classified as TMEM 0 and for those with homozygous C were classified as TMEM 1. One CTE-H sample was omitted from APOE/TMEM (missing TMEM106B), to keep samples constant between the risk variant study, the same sample was omitted from APOE analysis.

CTE-L and CTE-H samples were categorized based on the McKee staging criteria^13,14^ Stages I & II and Stages III & IV, respectively. RHI subjects were selected as athletes with a similar age at death as the CTE-L subjects and a similar level of repetitive head impacts exposure as the CTE subjects but exhibiting no CTE pathology (McKee Stage 0) upon autopsy. All subjects and brain samples were chosen as to avoid comorbidities and non-CTE pathology, including tau, as much as possible and had fresh frozen tissue available. All available brains possessed by the brain bank at the time of the study that matched these criteria were used. Total years of play was collected from family members at the time of brain donation and made available by the Brain Bank^18^. AT8 histology measurements were collected using digital pathology analyses via an Aperio slide scanner as previous described^19^. The raw AT8 measures were observed to be log-normal by inspection and therefore were log transformed for all downstream analyses. For this study, subjects with no APOE4 allele were classified as APOE 0, and those with at least one APOE4 allele were classified as APOE 1. Additionally, subjects with a G in TMEM106B rs3173615 were classified as TMEM 0 and those with homozygous C were classified as TMEM 1.

### 4.2 Sample processing, RNA Extraction, and mRNA Sequencing

Grey matter tissue was dissected from the cortical ribbon of the Brodmann Area 9 gyral crest to avoid sampling highly degenerated tissue. Total RNA was extracted using the Promega Maxwell RSC simplyRNA Tissue Kit (Cat No. AS1340) according to the manufacturer’s protocol. The integrity and quality of RNA were verified by an Agilent 2100 Bioanalyzer using RNA 600 Nano Chips (Cat No. 5067-1511). Only cases with a RNA Integrity Number (RIN) of 4 or higher using an Agilent Bioanalyzer instrument were selected for study. Paired end poly-A selected mRNA sequencing libraries with a targeted library size of 80M reads were generated using a Kapa Stranded mRNA-Seq Kit according to the manufacturer instructions and sequenced on an Illumina MiSeq instrument by the Boston University Microarray and Sequencing Core. Samples were sequenced in four batches due to study size and were distributed among batches as to avoid any confounding of status, age at death, RIN, or other important experimental variables.

### 4.3 mRNA-Seq Analysis

A custom analytical pipeline was developed to analyze the sequencing data. Sequencing reads adapter-and quality-trimmed using trimmomatic^20^ and assessed for high quality using FastQC^21^ and multiqc^22^. Trimmed reads were aligned against the human reference genome build GRCh38 with Gencode v34^23^ gene annotation using STAR^24^. Aligned reads were quantified to the gene level using the HTSeq package^?^ and the Gencode v34 gene annotation. For each analysis, genes with abundance estimates less than 0 in at least 50% samples within each group were filtered out.

Differential expression (DE) analyses were conducted separately for case status, total years of play, and AT8 with the DESeq2^25^ package. The six DE analyses are listed and summarized in Table 1. Group comparisons of CTE-L and CTE-H were performed separately with all RHI samples. For total years of play, AT8, APOE and TMEM106B, CTE-L and RHI samples were grouped together to increase sample size, as modeling continuous variables on the groups separately had insufficient statistical power to detect meaningful associations. CTE-H samples were analyzed as a group for these variables. Three samples (RHIN_0052, CTES_0064, CTES_0079) were filtered out of the years of play analysis due to extreme values (> 30 years) that drove spurious DE results. One sample, CTES_0068 was filtered out of the APOE and TMEM analysis due to missing TMEM106B value. All analyses included age at death, RIN, and sequencing batch in the model as covariates in addition to the variable of interest. Gene associations were considered significant if they attained a false discovery rate (FDR) of less than 0.1.

Differential expression results from each analysis were subject to preranked gene set enrichment against the Gene Ontology annotation^26^ curated in the c5 MsigDB gene set database^27^ using the fgsea^28^ package. GO categories were considered significant if they attained an FDR of less than 0.1. To aid in interpretation of GO terms, each enriched term was manually categorized by the investigators into one of a set of high level biological categories a priori. Categories include blood brain barrier (BBB), extracellular matrix (ECM)/membrane, cell cycle, cytoskeleton, development, immune/inflammation, ion homeostasis, metabolism/mitochondria, neuron, protein processing, signaling, transcription/translation, and an “other” category for terms that were not easily categorized. The immune/inflammation and neuron categories were further subcategorized due to the large number of enriched terms and to aid in further interpretation. All custom analysis was performed with python, R, snakemake^29^ and Jupyter Lab^?^ software.

### 4.4 qPCR Validation

Delta CT expression values for an a priori set of 33 genes known to be implicated in CTE (full gene list in Supplemental Table S2) in a set of 54 overlapping BA9 brain samples (37 CTE-H, 9 CTE-L, and 8 RHI) that overlapped with the samples presented in this study were analyzed for differential expression. Linear regression models were constructed for each gene modeling qPCR expression values as a function of each variable of interest separately adjusting for RIN and age at death, consistent with the DE models conducted with the mRNASeq data. Genes were marked as concordant between the qPCR and DE analyses if the direction of effect agreed (i.e. both either positive or negative log2 fold change), irrespective of significance in either dataset. Concordant/discordant genes were tabulated into confusion matrices for each analysis and Fisher’s exact test was applied to each to assess the likelihood the degree of concordance could occur by chance using a right-tailed p-value. Separately, Spearman correlation was computed for qPCR vs mRNASeq DE log2 fold change values, to assess overall agreement in ranked effect size.

### 4.5 Data and Code Availability

Raw and processed read data have been deposited into GEO under accessions GSE157330. All results, analysis, and figure code for this project are available on an Open Science Framework project located at https://osf.io/guepa/.

## Supporting information

mRNASeq DE gene information for qPCR validation

qPCR gene information

Gene Ontology categorization mapping

